# Deciphering HLA-I motifs across HLA peptidomes improves neo-antigen predictions and identifies allostery regulating HLA specificity

**DOI:** 10.1101/098780

**Authors:** Michal Bassani-Sternberg, Chloé Chong, Philippe Guillaume, Marthe Solleder, HuiSong Pak, Philippe O Gannon, Lana E Kandalaft, George Coukos, David Gfeller

## Abstract

The precise identification of Human Leukocyte Antigen class I (HLA-I) binding motifs plays a central role in our ability to understand and predict (neo-)antigen presentation in infectious diseases and cancer. Here, by exploiting co-occurrence of HLA-I alleles across ten newly generated as well as forty public HLA peptidomics datasets comprising more than 115,000 unique peptides, we show that we can rapidly and accurately identify many HLA-I binding motifs and map them to their corresponding alleles without any *a priori* knowledge of HLA-I binding specificity. Our approach recapitulates and refines known motifs for 43 of the most frequent alleles, uncovers new motifs for 9 alleles that up to now had less than five known ligands and provides a scalable framework to incorporate additional HLA peptidomics studies in the future. The refined motifs improve neo-antigen and cancer testis antigen predictions, indicating that unbiased HLA peptidomics data are ideal for *in silico* predictions of neo-antigens from tumor exome sequencing data. The new motifs further reveal allosteric modulation of the binding specificity of HLA-I alleles and we unravel the underlying mechanisms by protein structure analysis, mutagenesis and *in vitro* binding assays.

## Introduction

HLA-I molecules play a central role in defence mechanisms against pathogens and immune recognition of cancer cells. Their main functionality is to bind short peptides (mainly 9- to 12-mers) coming from degradation products of endogenous or viral proteins. The peptides are cleaved in the proteasome, transported by the transporter associated with antigen processing (TAP) complex, loaded onto the HLA-I molecules in the ER and presented at the cell surface [1]. Non-self peptides presented on HLA-I molecules, such as those derived from degradation of viral proteins, mutated proteins (referred to as neo-antigens), and other cancer specific and abnormally expressed proteins can then be recognized by CD8 T cells and elicit cytolytic activity. Neo-antigens have recently emerged as promising targets for development of personalized cancer immunotherapy [2].

Human cells express three HLA-I genes (HLA-A/B/C). These genes are the most polymorphic of the human genome and currently more than 12,000 different alleles have been observed [3]. Such a high polymorphism makes it challenging to model the different binding specificities of each allele and predict antigens presented at the cell surface. Information about binding motifs (mathematically defined here as Position Weight Matrices and graphically represented as sequence logos) of HLA-I molecules has been mainly obtained from biochemical assays where chemically synthesized peptides are tested *in vitro* for binding. This *in vitro* approach is experimentally laborious, time consuming and expensive. Currently, the most frequent HLA-I alleles have thousands of known ligands that provide a detailed description of their binding specificity. Many of these ligands are stored in very important resources such as IEDB [4,5] and have been used to train machine learning algorithms for HLA-I peptide interaction predictions [6-11]. However, the vast majority (>95%) of HLA-I alleles still lack documented ligands and despite very valuable algorithmic developments to generalize prediction methods to any allele [12], it remains more challenging to make accurate predictions for alleles without known ligands. Moreover, many alleles with known motifs are supported by only a few tens of peptides and some of theses ligands have been selected based on *a priori* expectations of the binding specificity rather than unbiased screening of random peptide libraries. Such potentially biased datasets can be sub-optimal for training HLA-I ligand predictors.

Mass-spectrometry (MS) analysis of HLA-I binding peptides eluted from cell lines or tissue samples is a promising alternative to the use of HLA-I ligand interaction predictions. Despite technical and logistic challenges (typically 1cm^3^ of tissue material is required or 1x10^8^ cells in culture), MS is increasingly used to directly identify viral [13,14] or cancer-specific (neo-)antigens [15-20]. While MS is currently the most promising tool for the discovery of the *in vivo* presented neo-antigen, it is only applicable to a small fraction of samples due to the large amount of material that is required and the complexity of these experiments. However, tens of thousands of endogenous peptides naturally presented on HLA-I molecules are identified in such HLA peptidomics studies, providing a unique opportunity to collect very large numbers of HLA-I ligands that can be used to better understand the binding properties of HLA-I molecules. The challenge in studying HLA-I motifs based on such pooled peptidomics data from unmodified cell lines or tissue samples is to determine the allele on which each peptide was displayed. The most widely used approach is to predict binding affinity of each peptide to each allele present in a sample [21]. Recent studies have suggested that HLA-I motifs could be identified in HLA peptidomics datasets in an unsupervised way by grouping peptides based on their sequence similarity [17,22-24]. However, this strategy still relies on previous information about HLA-I binding specificity when associating predicted motifs with HLA-I alleles and is therefore restricted to alleles whose motifs have been already characterized.

Here, we describe a computational framework for direct identification of dozens of HLA-I motifs without any *a priori* information about HLA-I binding specificity by taking advantage of co-occurrence of HLA-I alleles across both newly generated and publicly available HLA peptidomics datasets. Our approach recapitulates and refines motifs for many common alleles and uncovers new motifs for eight alleles for which, until this study, no ligand had been documented. Importantly, this approach is highly scalable and will enable continuous refinement of motifs for known alleles and determination of novel motifs for uncharacterized alleles as more HLA peptidomics data will be acquired in the future. Training HLA-I ligand predictors based on the refined motifs significantly improves neo-antigen predictions in five out of six tumor samples with experimentally determined neo-antigens. Our large collection of HLA-I ligands further allowed us to unravel some of the molecular determinants of HLA-I binding motifs and revealed allosteric modulation of HLA-I binding specificity. To elucidate the underlying molecular mechanisms, we show how a single point mutation in HLA-B14:02 outside of the B pocket significantly changes the amino acid preferences at P2 in the ligands.

## Results

### Fully unsupervised identification of HLA-I binding motifs

To study the binding properties of HLA-I alleles without relying on *a priori* assumption on their binding specificity and investigate whether this unbiased approach could improve neo-antigen predictions from exome sequencing data, we reasoned that HLA-I binding motifs might be identified across samples with in-depth and accurate HLA peptidomics data by taking advantage of co-occurrence of HLA-I alleles. To this end, we measured the HLA peptidome eluted from six B cell lines, two *in vitro* expanded tumor-infiltrating lymphocytes (TILs) samples and two leukapheresis samples (peripheral blood mononuclear cells) selected based on their high diversity of HLA-I alleles (see Methods and S1 Dataset). By applying a stringent false discovery rate for peptides identification of 1%, we accurately identified 47,023 unique peptides displayed on 32 HLA-I molecules. To expand the coverage of HLA-I alleles, we further collected 40 publicly available high-quality HLA peptidomics datasets [17,18,22,23,25-27] (see Methods and S2 Dataset). Our final data consists of a total of 50 HLA peptidomics datasets covering 66 different HLA-I alleles (18 HLA-A, 32 HLA-B and 16 HLA-C alleles, see Supplementary Table 1). The number of unique HLA-I ligand interactions across all samples reaches 252,165 for a total of 119,035 unique peptides (9-to 14-mers), which makes it, to our knowledge, the largest currently available collection of HLA peptidomics datasets both in terms of number of peptides and diversity of HLA-I molecules. Binding motifs in each HLA peptidomics dataset were identified for 9-and 10-mers using a motif discovery algorithm initially developed for multiple specificity analyses [28,29] and recently applied to the analysis of small (seven) HLA peptidomics datasets [24] (see S1 Fig). Importantly, this method does not rely on HLA-I peptide interaction predictions (see Methods).

**Table 1:**
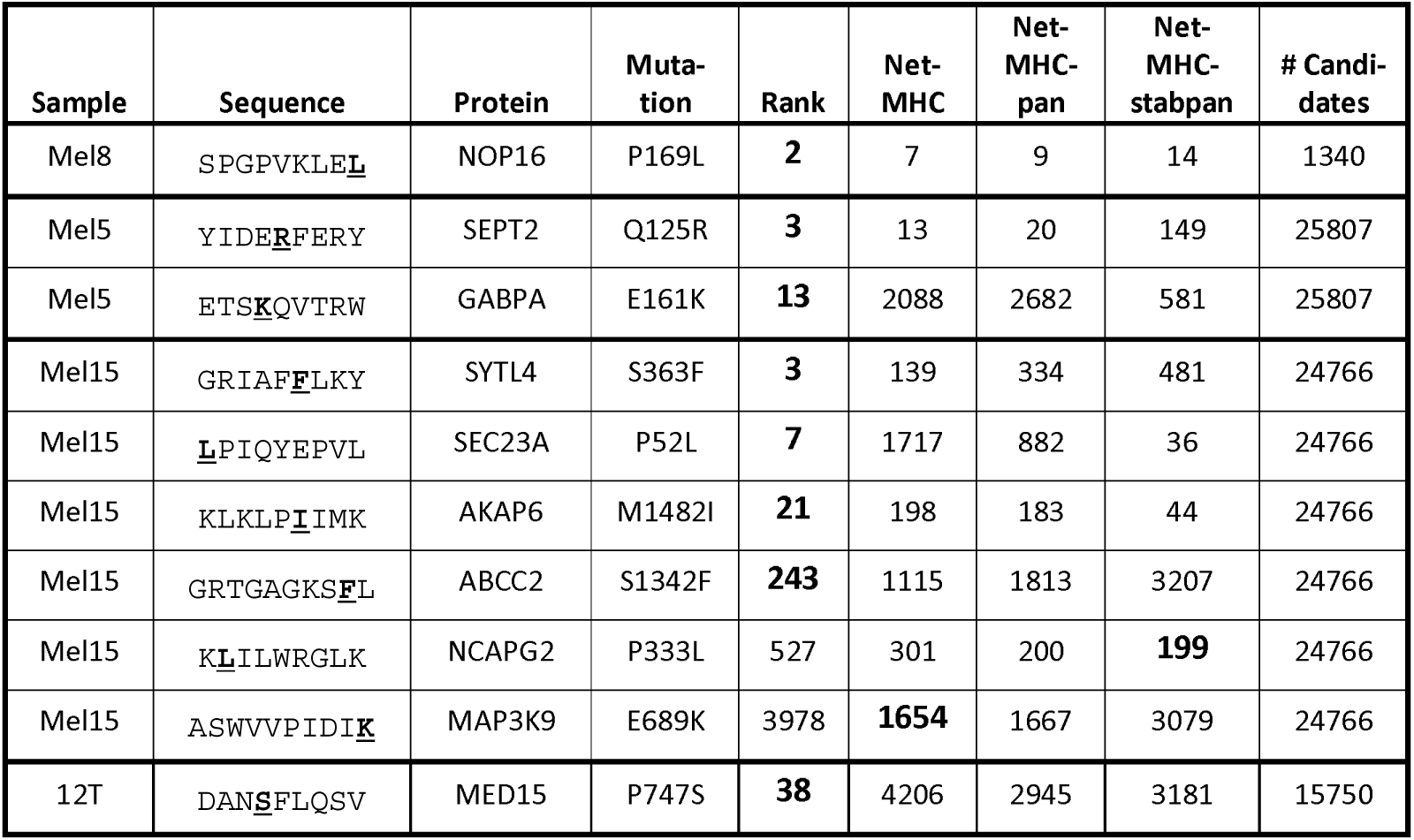
Ranking of the neo-antigens identified in four melanoma samples [17,20]. Column 2 shows the mutated neo-antigens (the mutated residue is highlighted in bold). Column 5 shows the ranking based on our predictions. Columns 6 to 8 show the ranking based on NetMHC [8], NetMHCpan [12] and NetMHCstabpan [34], respectively. The last column shows the total number of neo-antigen candidates (i.e., all possible 9-and 10-mers encompassing all missense mutations).

To assign each motif to its allele even in the absence of *a priori* information about the alleles’ binding specificity, we developed a novel computational strategy illustrated in Fig. 1. In this example, one allele (HLA-A24:02) was shared between all three samples. Remarkably, exactly one identical motif was shared between the three samples. As such, one can predict that this motif corresponds to the shared allele. Similarly, two alleles (A01:01 and C06:02) were shared between exactly two samples and here again two motifs were shared among the corresponding samples, and could therefore be annotated to their corresponding alleles. Moreover, if one sample shares all but one allele with another sample, it can be inferred that the motif that is not shared corresponds to the unshared allele (see example in S2 Fig). Finally, if all but one motif had been annotated in a sample to all but one allele, one can infer that the remaining motif corresponds to the remaining allele. These three ideas can then be recursively applied to identify HLA-I motifs across our large collection HLA peptidomics datasets (see Methods). Of note, motifs identified in distinct samples that have some alleles in common show very high similarity (Fig 1 and S2 Fig) and our new approach builds upon this remarkable inherent reproducibility of in-depth and accurate HLA peptidomics data.

**Fig 1:**
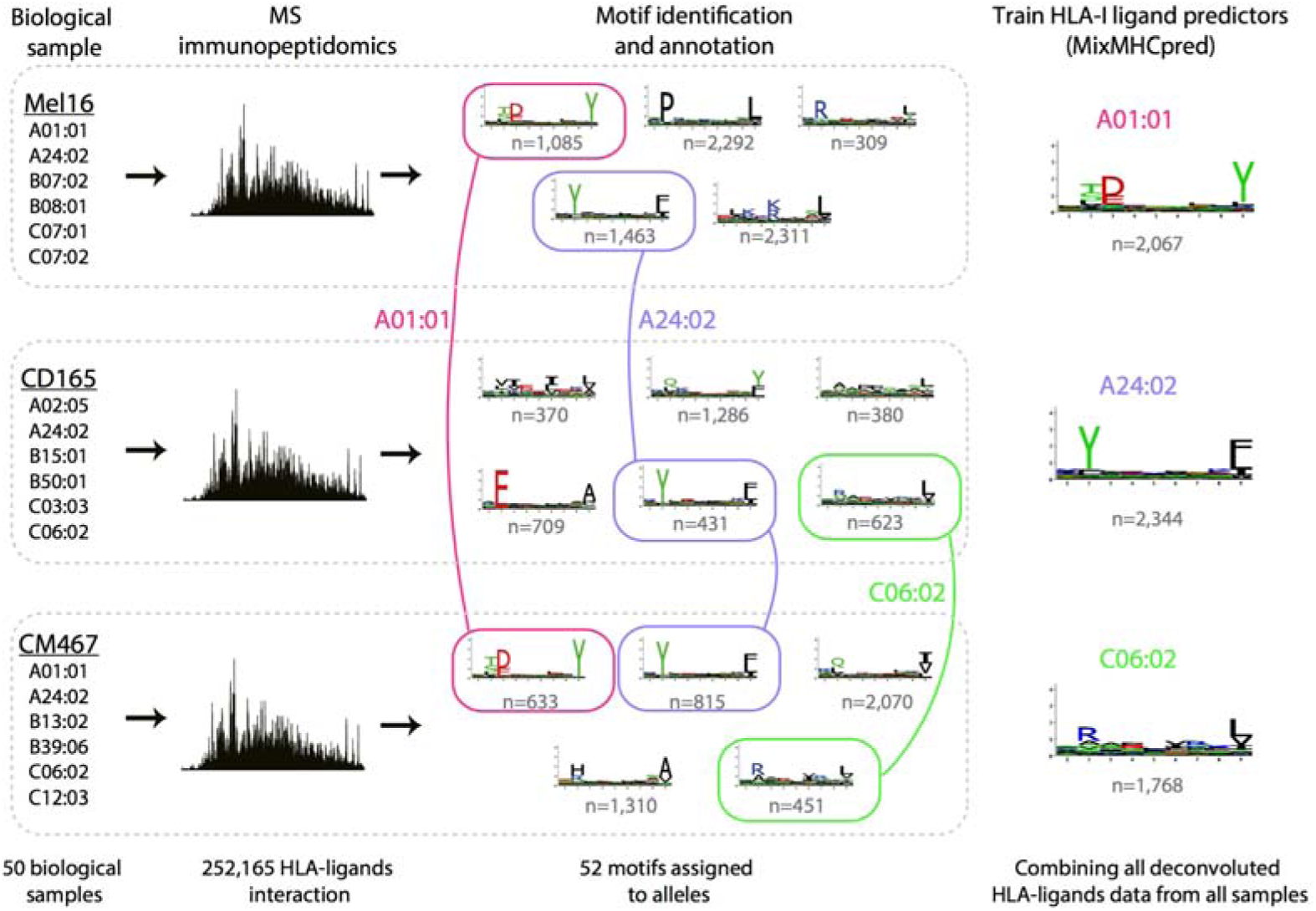
General pipeline for HLA-I motif identification and annotation, and training of predictors. High accuracy HLA peptidomics data were first generated for 10 samples and collected from publicly available data for 40 other samples. In each sample motifs were identified using on our recent mixture model algorithm [24]. Motifs were then annotated to their respective allele based on co-occurrence of alleles across samples (e.g., first HLA-A24:02, then HLA-A01:01 and HLA-C06:02, see also S2 Fig for another example). Finally all peptides assigned to each motif were pooled together to train our new HLA-I ligand predictor (“MixMHCpred") for each HLA-I allele.

We applied our algorithm to the 50 HLA peptidomics datasets considered in this study. In total, 44 different motifs could be associated with a specific allele without relying on any *a priori* assumption of HLA-I binding specificity (Fig. 2A). These include seven alleles (HLA-B13:02, HLA-B14:01, HLA-B15:11, HLA-B15:18, HLA-B18:03, HLA-B39:24 and HLA-C07:04) that did not have known ligands in IEDB, and 5 additional ones (HLA-B38:01, HLA-B39:06, HLA-B41:01, HLA-B56:01, HLA-C07:01) that had less than 50 known ligands. To validate our predictions, we compared the motifs predicted by our fully unsupervised method with known motifs derived from IEDB [4], when available. Despite some differences (e.g. HLA-A25:01 motif at P9) affecting especially alleles with low number of ligands in IEDB (S3 Fig), we observed an overall high similarity confirming the reliability of our predicted motifs (Fig. 2A). However, it is important to realize that even small differences in the motifs can have important effects on the performance of predictors that are trained on such data when ranking very large lists of potential epitopes. When comparing with data recently obtained by HLA peptidomics analysis of mono-allelic cell lines [30], a very high similarity was also observed (Fig. 2B, stars in S3 Fig), which further validates our computational approach for HLA-I motif identification and annotation from in-depth pooled HLA peptidomics data.

**Fig 2:**
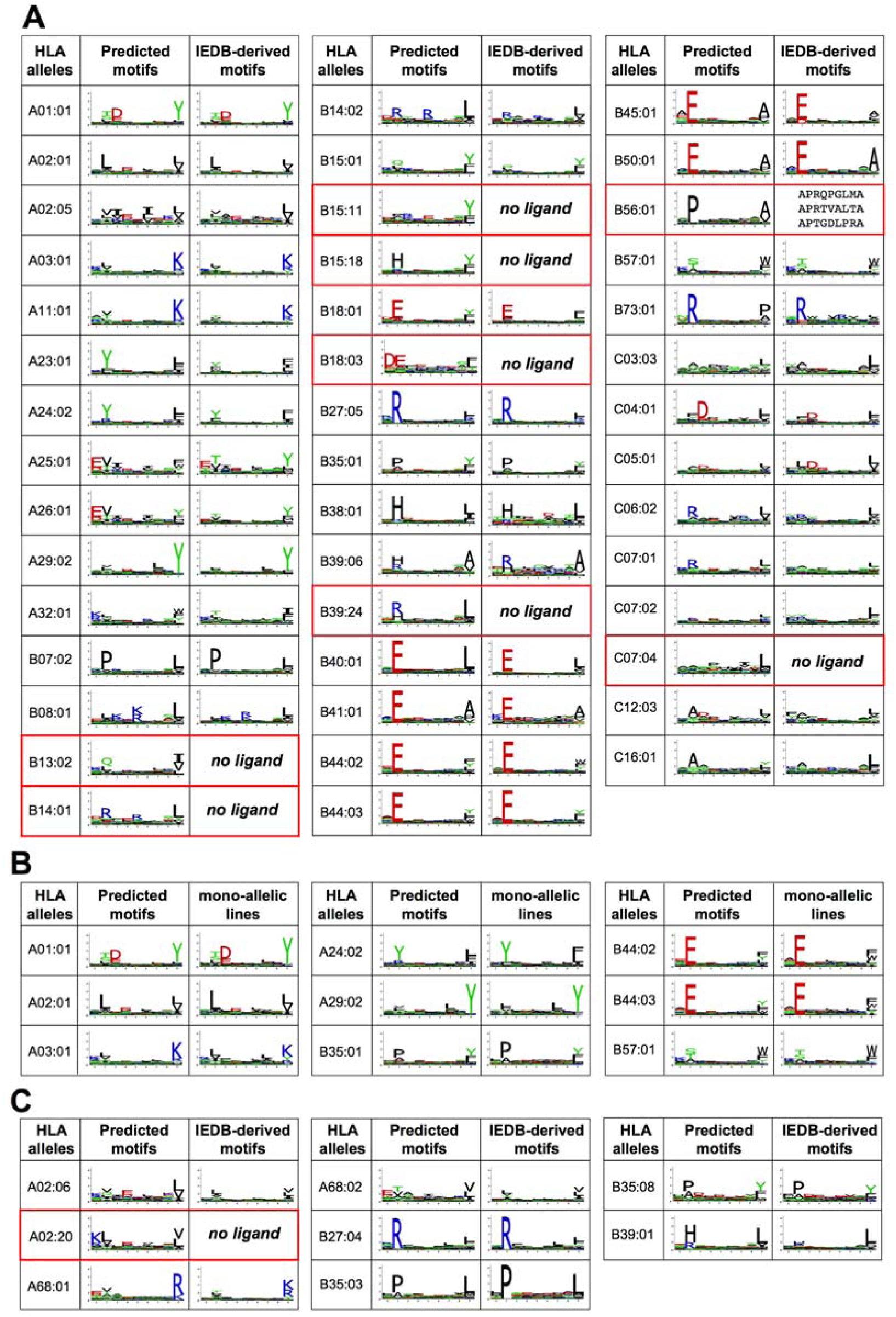
Comparison between motifs predicted by our algorithm and known motifs. **A**: Comparison with IEDB motifs for 44 HLA-I binding motifs identified with the fully unsupervised approach. Alleles without previously documented ligands are highlighted in red. For HLA-B56:01, the three known ligands are shown. **B**: Comparison with motifs obtained from mono-allelic cell lines [30]. **C**: Motif identified with the semi-supervised approach.

As expected from many previous studies, alleles with the same two first digits code showed high similarity in their binding specificity, apart from HLA-B15 alleles which are known to be more diverse [31]. This includes many of the new motifs (e.g., HLA-B14:01 vs HLA-B14:02; HLA-B18:03 vs HLA-B18:01; HLA-B39:24 vs HLA-B39:06), which provides further evidences of the accuracy of our predictions for these uncharacterized alleles.

### Semi-supervised approach

For the most frequent HLA-I alleles, including several shown in Fig. 2A, a good description of their binding motifs can be already obtained from existing databases [4]. To further expand our collection of HLA-I binding motifs, we used similarity to the binding motifs derived from IEDB ligands to annotate motifs that could not be assigned to their corresponding allele by the fully unsupervised approach (see Methods and [24]). This comparison typically included one or two clearly distinct motifs that had not been annotated by the fully unsupervised approach and it could be reliably performed in most samples. This enabled us to determine the binding motifs of 8 additional alleles (Fig. 2C). Of note, the new motif of HLA-A02:20 was predicted by observing that it was the only motif not annotated in one sample (RA957) and could only be annotated to this allele based on the motifs identified in other samples for all the other alleles (see S1 Fig). The final list of motifs for the 52 alleles and detailed comparison with IEDB derived data, when available, is shown in S4 Fig. Importantly, for the majority of alleles considered in this study, the motifs are supported by significantly more ligands than what is available in existing databases (S5 Fig).

The full pipeline was also applied on 10-mers identified by MS across the 50 HLA peptidomics studies and revealed six new motifs for poorly characterized alleles in IEDB (S6 Fig).

### Investigating inherent technical biases in HLA peptidomics studies

Different technical biases may affect MS data, which could undermine their use for training HLA-I ligand predictors. To investigate this potential issue, we computed amino acid frequencies at non-anchor positions (P4 to P7) in our HLA peptidomics data, excluding alleles displaying anchor residues at these positions (see Methods and Supplementary Table 2). The reason for focusing on middle positions is that they display low specificity (especially in 9-mers, see discussion in [24,32,33] for longer peptides) and therefore could provide a global view of potential MS biases on amino acid frequencies that is not affected by the constraints of binding to HLA-I molecules. As expected, we observed a high correlation between amino acid frequencies at non-anchor positions in our HLA peptidomics data and in the human proteome (r=0.85) (Fig. 3 and S7 Fig). The most important difference was found for cysteine, which is prone to post-translational modifications that are typically not included in database searches and was observed at very low frequency in the HLA peptidomics data (the same observation was recently made in mono-allelic cell lines [30]). Other amino acids were not much under-or over-represented, and no clear pattern emerged from these data with respect to amino acid biophysical properties (e.g., charge, hydrophobicity, size). Overall, our results suggest that HLA peptidomics data do not show strong technical biases, apart from under-representation of cysteine which can be compensated by re-normalisation (see next section), and therefore could provide ideal data for training HLA-I peptide interaction predictors, especially for ligands coming from human cells like neo-antigens.

**Fig 3:**
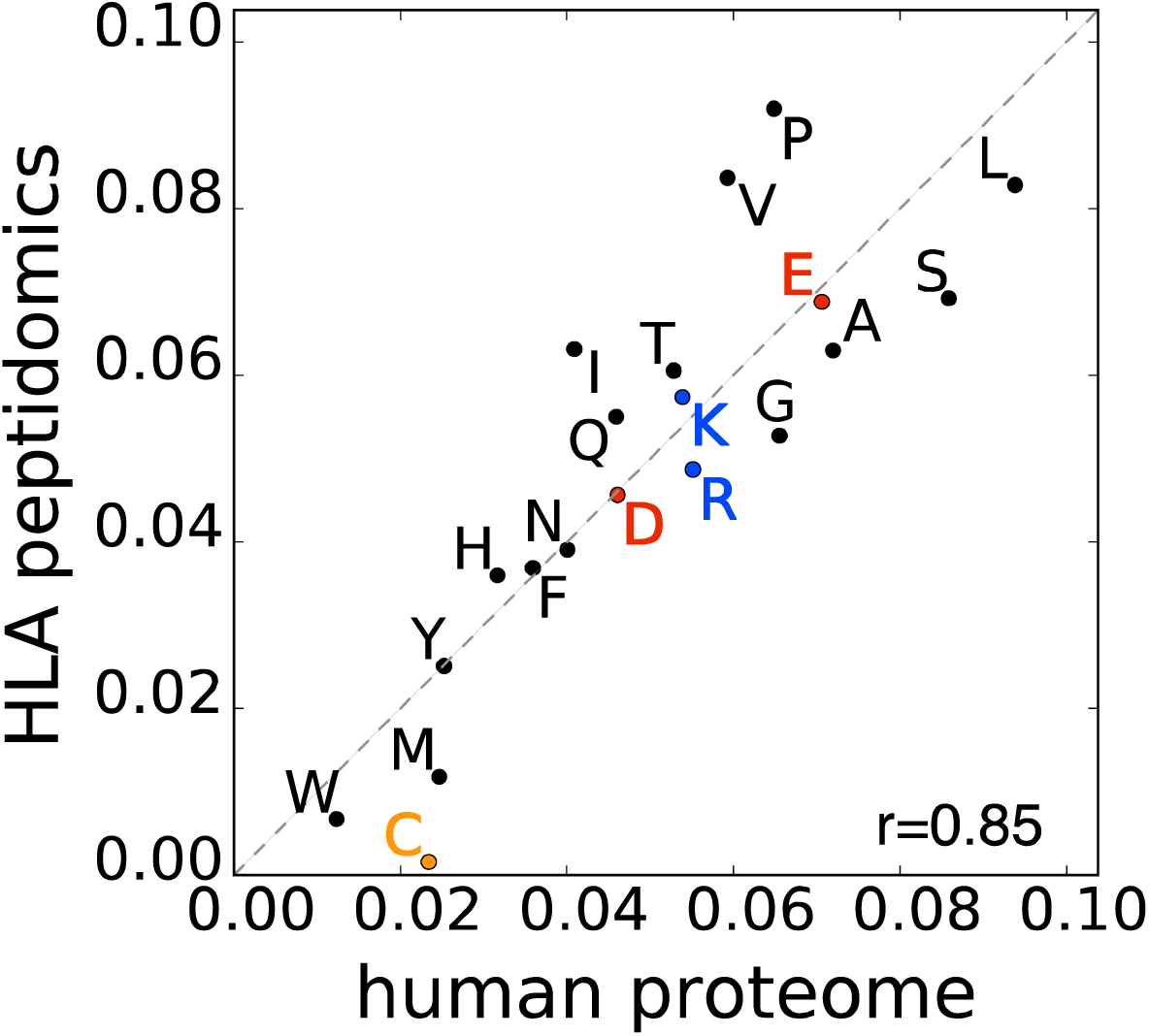
Correlation between amino acids frequencies at positions P4 to P7 in our HLA peptidomics data and in the human proteome.

### Training HLA-I ligand predictors

To test whether our unique dataset of naturally presented peptides could help predicting HLA-I ligands, including neo-antigens in tumors based on exome sequencing data, we trained a predictor of HLA-I ligands (referred to as “MixMHCpred"). As MS only includes positive examples and HLA-I ligands in general do not show strong amino acid correlations between different positions (see discussion in [24] for some exceptions), we built Position Weight Matrices (PWMs) for each of the 52 alleles. These PWMs were built by pooling together all peptides assigned to each allele across all our HLA peptidomics datasets (see Methods and Fig. 1). We further included MS data from mono-allelic cell lines for 6 rare alleles that were not present in our dataset, resulting in a total of 58 alleles available in our predictor. To correct for the low detection of cysteine observed in HLA peptidomics data we further renormalized our predictions by amino acid frequencies at non-anchor positions (see Methods).

As a first validation, we attempted to re-predict naturally presented peptides experimentally identified in ten mono-allelic cell lines whose alleles overlapped with our dataset [30]. For this analysis, we did not include data from these mono-allelic cell lines in our training set. To assess our ability to predict naturally presented peptides, we added 99-fold excess of decoy peptides randomly selected from the human proteome to each mono-allelic cell line dataset and measured the fraction of Positives among the top 1% Predictions (PP1). For all but one allele, our algorithm showed higher predictive power compared to standard HLA-I ligand predictors [8,12,34] (Fig 4A).

**Fig 4:**
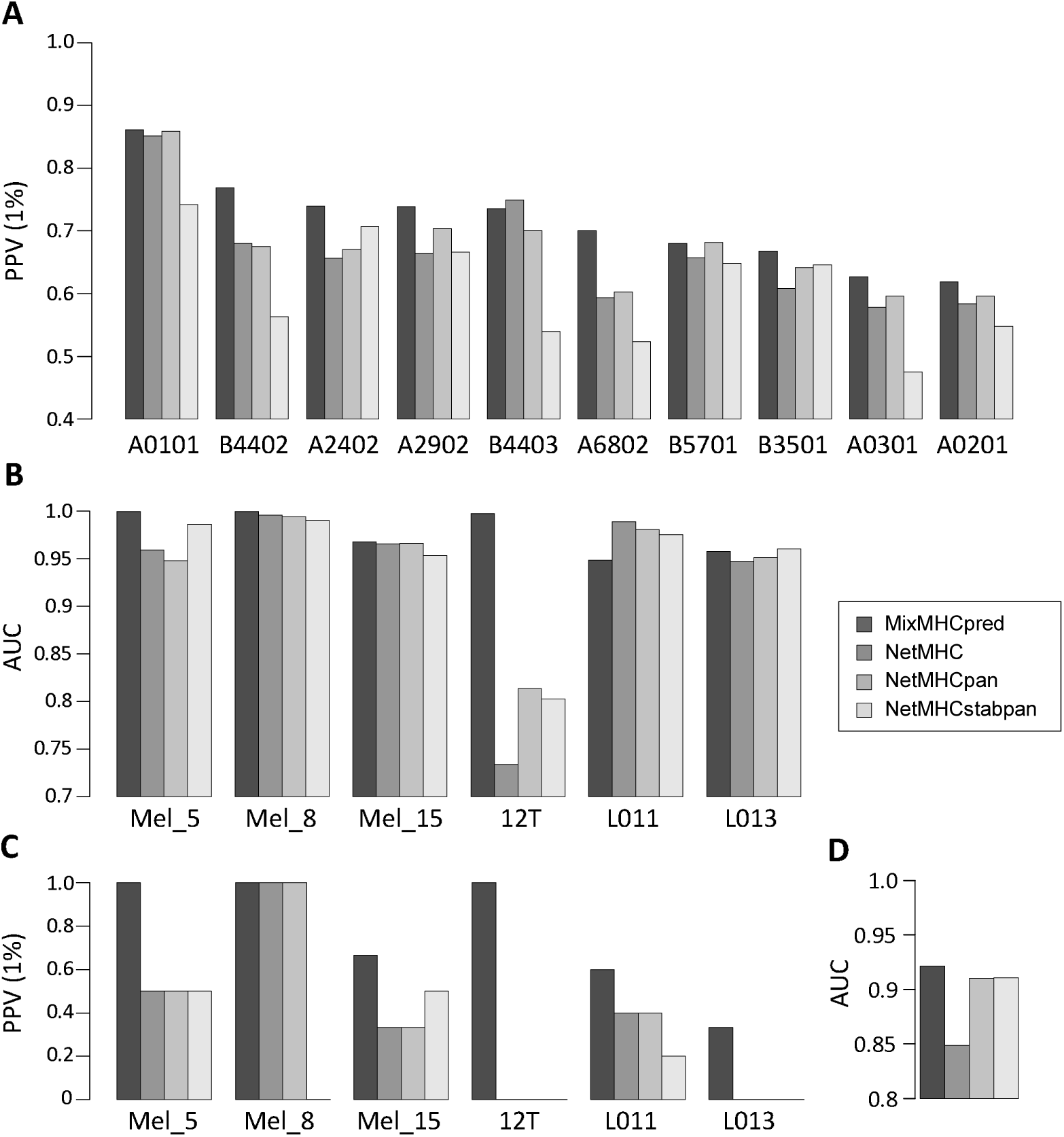
Comparison between our predictor (MixMHCpred) and existing tools. **A**: Fraction of true positives among the top 1% predictions for the naturally presented HLA-I ligand identified in mono-allelic cell lines, with 99-fold excess of decoy peptides (in this case, all AUC values were all larger than 0.99, which makes it difficult to use them to compare between methods). **B:** AUC values for neo-antigen predictions. In Mel_5, Mel_8, Mel_15 and 12T the neo-antigens were identified without relying on HLA-I ligand predictions. In L011 and L013, HLA-I ligand predictions were used to pre-select the peptide to be tested experimentally. **C**: Fraction of neo-antigens among the top 1% predictions (same samples as in B). **D:** AUC values obtained for cancer testis antigens from the CTDatabase.

### Predictions of neo-antigens and cancer testis antigens

We then collected currently available datasets that included direct identification of neo-antigens displayed on cancer cells as well as exome sequencing (Mel5, Mel8, Mel15 from [17] and 12T from [20]) for a total of ten 9-and 10-mers mutated peptides experimentally found to be presented on cancer cells (see Table 1). This dataset has the unique advantage of not being restricted to peptides selected based on *in silico* predictions, and is therefore an ideal testing set for benchmarking our predictor. Moreover, as these studies are quite recent, the neo-antigens used here as testing set are not part of the training set of any existing algorithm. In particular, they are not part the large training set used in this study since we only included wild-type human peptides in our pipeline. We then retrieved all possible 9-and 10-mer peptides that encompassed each missense mutation (S3 Dataset) and ranked separately for each patient these potential neo-antigens based on the score of our predictor (see Methods and Table 1). Remarkably, six of the ten neo-antigens fell among the top 25 predicted peptides, suggesting that by testing as few as 25 mutated peptides per sample, we could identify more than half of the neo-antigens identified by MS (Table 1). Considering that the total number of potential neo-antigens (i.e. 9-and 10-mers containing a missense mutation) can be as large as 25,000 for tumors with high mutational load, our predictor trained on naturally presented human HLA-I ligands clearly enabled us to significantly reduce the number of peptides that would need to be experimentally tested to identify *bona fide* neo-antigens from exome sequencing data. We further added two datasets of neo-antigens identified in lung cancer patients in a recent study (L011 and L013) [35], although only peptides pre-selected based on binding affinities predicted with existing tools [12] were tested in this study. Here again, our predictor ranked one neo-antigen in the top 25 predicted peptides in both samples (Supplementary Table 3). When comparing with standard tools that are widely used to narrow-down the list of potential neo-antigens predicted from exome sequencing data [8,12,34], our method trained on HLA peptidomics data showed clear improvement with a mean AUC value of 0.979, compared to 0.932 for NetMHC [8], 0.942 for NetMHCpan [12] and 0.945 for NetMHCstabpan [34] (Fig 4B) and increased number of neo-antigens in the top 1% predictions across all six samples (Fig 4C). Importantly, even if we did not include in the training of our predictor MS data (i.e. wild-type peptides) from the samples in which the neo-antigens were identified, neo-antigens were still more accurately predicted compared to other tools (see S8 Fig). This demonstrates that our approach for neo-antigen predictions from the list of somatic mutations identified by exome sequencing of tumors does not require HLA peptidomics data from the same sample where neo-antigens had been identified. Nevertheless, both our predictor and standard prediction tools failed to identify some neo-antigens (e.g., KLILWRGLK from NCAPG2 P333L mutation, see Table 1). This suggests that, when enough tumor material is available for immunopeptidomics analyses, direct identification of neo-antigens with MS should still be performed to optimally enrich in true positives the list of ligands to be experimentally tested for immunogenicity [15,19].

The number of studies reporting both neo-antigens and exome sequencing results is still limited. To benchmark our algorithm with larger datasets of immunologically relevant tumor antigens, we tested our ability to predict epitopes from cancer testis antigens. We retrieved epitopes listed in the CTdatabase [36] (see Methods and Supplementary Table 4). We then assessed how our predictor could prioritize these epitopes from all possible peptides encoded by these cancer testis antigens. Although we cannot exclude that some of these epitopes had been selected for experimental testing after prediction by older versions of HLA-I ligand predictors, we still observed improvement using our predictor trained only on naturally presented HLA-I ligands (Fig 4D). This indicates that improvement in prediction accuracy is not restricted to elution data (see similar results in [24]).

MS data can contain false positives for many different reasons, such as errors in the computational identification of peptides from the spectra or co-eluting contaminating peptides that bind unspecifically to the affinity column. Therefore, despite the high quality of HLA peptidomics datasets generated in this study (<1% FDR), we do expect our data to contain a few contaminants. To test the robustness of our motif discovery and annotation pipeline, and our HLA-I ligand predictor, we incorporated 5% of random peptides from the human proteome into all HLA peptidomics datasets considered in this work and rerun the whole motif annotation pipeline and training of the predictor. Remarkably, the accuracy of the predictions was only very modestly affected by this noise and predictions were still better than with other existing tools (S9 Fig). This suggests that our pipeline is robust to experimental contaminations in the data and indicates that the wealth of unbiased and accurate data provided by MS can compensate the inherent contaminations, when using these data for training HLA-I ligand predictors.

### Analysis of the newly identified motifs

One of our novel HLA-I motifs describes the binding specificity of HLA-A02:20 (Fig. 2C). HLA-A02 binding motifs have been widely studied. However, HLA-A02:20 motif differs from standard HLA-A02 motifs at P1, with a clear preference for charged residues (Fig. 5A). Interestingly, HLA-A02:20 is among the very few (<2%) HLA-A02 alleles that do not have a conserved lysine pointing towards P1 at position 90. Instead an asparagine is found there (Fig. 5A), and this residue is the only difference with the sequence of the very common A02:01 allele. To explore how the absence of lysine at position 90 could explain the observed difference in binding specificity, we collected all HLA-I alleles showing preference for charged amino acids at P1 (see S10 Fig). All of them had either asparagine or isoleucine at position 90. We then explored available crystal structures of HLA-I peptide complexes with charged residues at P1. HLA-B57:03 was crystalized with such a ligand (KAFSPEVI) [37]. Superposing the crystal structure of this complex with the structure of HLA-A02:01 provides a possible mechanism for understanding the change in binding specificity at P1. In HLA-A02:01, lysine at position 90 interacts with the hydroxyl group of serine at P1 (Fig. 5A, green sidechains). Such a conformation would not be compatible with a longer residue. Reversely, when asparagine was found at position 90, it did not point towards P1 (Fig. 5A, pink sidechains), thereby freeing space for larger sidechains like lysine or arginine at P1. Overall, our analysis indicates that the presence of asparagine at residue 90 may be responsible for the change in binding specificity between HLA-A02:01 and HLA-A02:20. More generally, our results suggest that lysine at residue 90 in HLA-I alleles strongly disfavours charged residues at P1.

**Fig 5:**
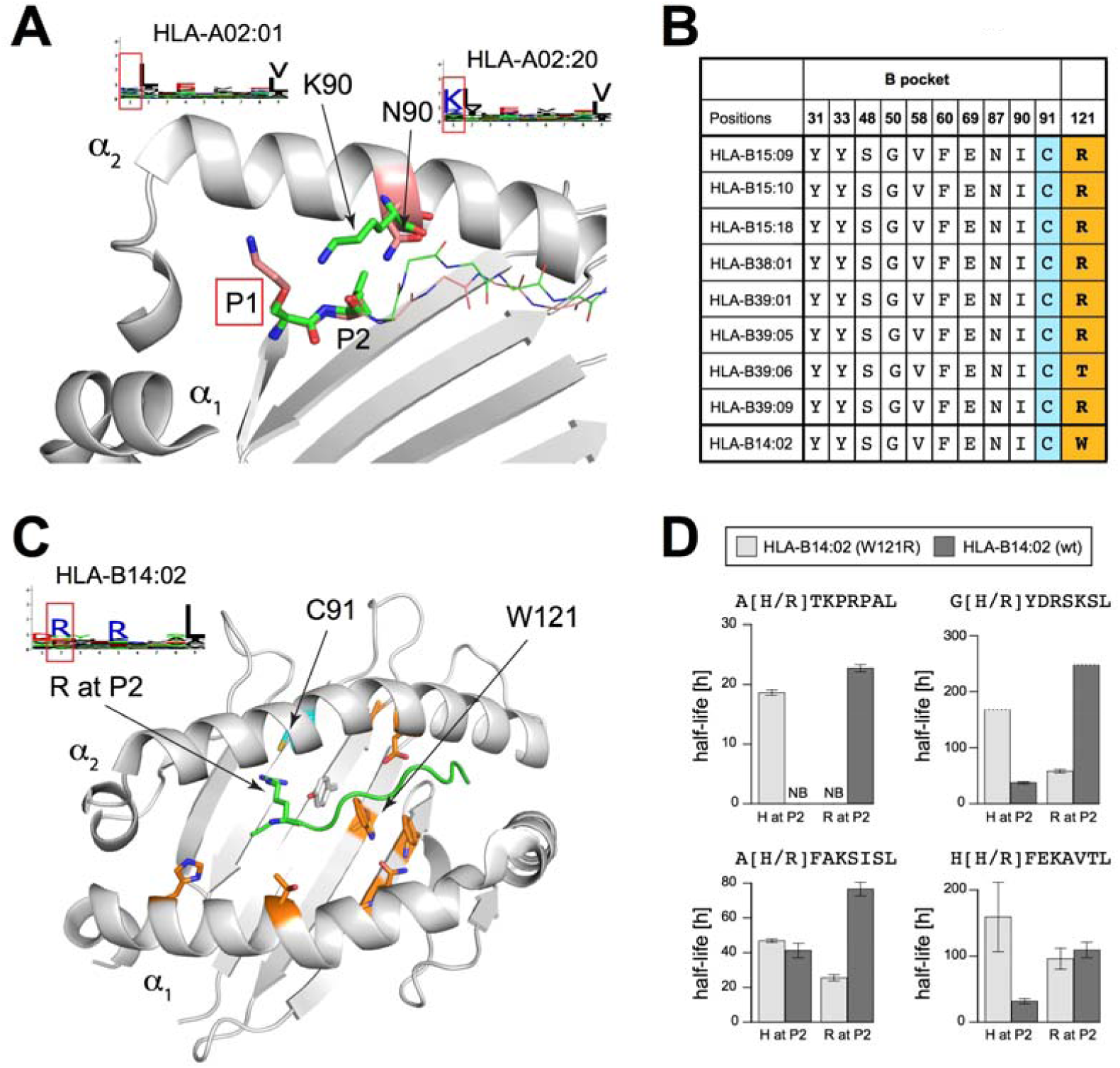
Analysis of newly identified HLA-I motifs. **A**: Structural view of two different HLA-I alleles with N90 as in HLA-A02:20 (PDB: 2BVQ [37], pink sidechains) or K90 as in HLA-A02:01 (PDB: 2BNR [52], green sidechains). For clarity, the α_1_ helix has been truncated. **B:** B pocket residues’ conservation across HLA-I alleles displaying preference for histidine at P2. The last line shows the sequence of HLA-B14:02, which does not show histidine preference at P2 (see motif in **C**), but has the same B pocket as HLA-B15:18. The last column shows amino acids at position 121, which is not part of the B pocket. **C:** Structural view of HLA-B14:02 in complex with a peptide with arginine at P2 (PDB: 3BVN [51]). Residues not conserved between HLA-B15:18 and HLA-B14:02 are displayed in orange. None of them are making direct contact with the arginine residue at P2. **D:** Stability values (half-lives) obtained for peptides with H or R at P2 for both HLA-B14:02 wt and W121R mutant. NB stands for no binding. Dashed lines indicate lower bounds for half-lives values.

The new motif identified for HLA-B15:18 (Fig. 2A) displayed strong preference for histidine at P2, which is not often observed in HLA-I ligands. To gain insights into the mechanisms underlying this less common binding motif, we surveyed all alleles that show preference for histidine at P2 (S11 Fig). Sequence and structure analysis showed that all of them have a conserved P2 binding site, commonly referred to as the B pocket (see Fig. 5B). However, several HLA-B14 alleles have exactly the same B pocket but show specificity for arginine at P2 (S11 Fig). Among them, HLA-B14:02 had the highest sequence similarity to HLA-B15:18, with only 8 different residues in the peptide binding domain, none of them making any contact with arginine at P2 in the crystal structure of HLA-B14:02 (orange residues in Fig. 5C). This suggests that the difference in binding specificity at P2 between HLA-B14:02 and HLA-B15:18 is likely explained by allosteric mechanisms. Of particular interest is residue 121 (W in HLA-B14:02 and R in HLA-B15:18), which is more than 7Å away from the arginine sidechain at P2 and is part of a network of aligned aromatic residues (Y33, W121 and F140) in HLA-B14:02 (Fig. 5C). We hypothesized that mutating this residue into arginine in HLA-B14:02 may modify the binding specificity to accommodate histidine at P2.

### Molecular mechanism underlying allosteric modulation of HLA-I binding specificity

To test our hypothesis, we generated a construct for HLA-B14:02 wild-type (wt) and W121R mutant. We tested several ligands of HLA-B15:18 identified in our HLA peptidomics data with histidine at P2, which were predicted to show enhanced binding to HLA-B14:02 W121R. As expected, a strong decrease in binding stability was observed between HLA-B14:02 W121R and HLA-B14:02 wt (Fig. 5D). Reversely, when testing the same peptides with arginine at P2, a significant increase in stability was observed between HLA-B14:02 W121R and HLA-B14:02 wt (Fig. 5D). For instance, binding of the peptide AHTKPRPAL was fully abolished in HLA-B14:02 wt, but was rescued when changing histidine to arginine at P2. Although other residues may also play a role in the binding specificity differences between HLA-B14:02 and HLA-B15:18, all of them are further away from P2, which supports the allosteric hypothesis. Overall, our results show that HLA-I binding specificity at P2 can be modulated by amino acids outside of the B pocket, and these binding experiments further confirm the motifs predicted for HLA-B14 alleles.

## Discussion

Despite decades of work to characterize the binding motifs of the most common HLA-I alleles, unbiased peptide screening approaches have not been commonly used in the past. This is mainly because both the N-and the C-terminus of the peptides are engaged in binding to HLA-I molecules, thereby preventing the use of high-throughput techniques for peptide screening like phage display. To address this issue, we developed novel algorithms to rapidly identify and annotate HLA-I binding motifs in a fully unsupervised way using in-depth HLA peptidomics data from unmodified cell lines and tissue samples. This enabled us to refine models of binding specificity for many alleles with few ligands in existing databases and characterize the binding properties of eight HLA-I alleles that had no known ligands until this study. Our approach is conceptually similar to existing approaches to deconvolute peptide epitopes by identifying shared peptides between different pools showing T cell reactivity in ELISpot experiments, but had never been applied to motif annotation across HLA peptidomics datasets.

Remarkably, our predicted motifs displayed high similarity with known motifs for common alleles, including motifs derived from HLA peptidomics analyses of mono-allelic cell lines [30] (Fig 2), and the MS-induced technical bias (mainly low detection of cysteine) could be compensated by normalization with expected amino acid frequencies. This suggests that HLA peptidomics data are optimal to train HLA-I ligand interaction predictors, as confirmed by our ability to accurately predict from exome sequencing data several neo-antigens identified in tumor samples. These observations are in line with recent results obtained with predictors trained on HLA peptidomics data from mono-allelic cell lines for 16 human class I alleles [30] and mouse class II alleles [38]. Although mass spectrometry may miss a significant fraction of the actually presented and immunogenic neo-antigens, those detected by MS are likely presented at high level on cancer cells. Therefore, accurately predicting such dominant neo-antigens is promising to prioritize targets for cancer immunotherapy.

Importantly, the improvement in prediction accuracy we report here comes primarily from refinement of known HLA-I motifs, since the less frequent alleles for which we uncovered new motifs were in general not part of our testing sets. For instance, differences are observed at P9 between the motif of HLA-A25:01 obtained from HLA peptidomics data (preference for F/W/Y/L) and from IEDB ligands (preference for Y/L/M/F) (Fig. 2A). This likely explains why the neo-antigen ETS**K**QVTRW was poorly predicted by standard tools [8,12] (predicted IC_50_ > 3,000nM). Although these differences may look relatively small on the logos (Fig 2A), they play an important role when ranking tens of thousands of potential epitopes with HLA-I ligand predictors. Moreover, as our predictor is only trained on naturally presented ligands, it may also capture some features of antigen presentation of endogenous peptides beyond the binding to HLA-I molecules. Along this line, it is interesting to note that a less important improvement in predictions had been observed when attempting to predict all ligands from the SYFPEITHI database [39] (including a large fraction of viral peptides) with HLA peptidomics data [24]. Although the dataset used in this previous study was significantly smaller, this observation suggests that HLA peptidomics data may be especially well suited for training predictors of human endogenous or mutated HLA-I ligands. Overall, our work highlights the importance of carefully determining HLA-I motifs, including for alleles that already have some known ligands, based on unsupervised analysis of naturally presented human HLA-I ligands for neo-antigen discovery.

Currently our predictor is limited to 9-and 10-mers, which is the most common length of HLA-I ligands and accounts for more than 80% of the HLA peptidome [17]. Although motif identification may work in some cases for 11-mers [24], the automated motif deconvolution and annotation becomes less accurate, especially for samples with less than 10’000 peptides identified by MS. Therefore, rather than including sub-optimal motifs in our predictor, we focused in this work on 9-and 10-mers. We anticipate that manual curation of HLA-I motifs in pooled HLA peptidomics datasets or the use mono-allelic cell lines [30] may be more appropriate for training predictors for longer peptides. Importantly, not including 11-mers has no influence on the predictions for 9-and 10-mers, since peptides of different length are treated separately in the current framework (see ref. [8] for possible algorithmic extensions to include peptides of different length in the training set of HLA-I ligand predictors).

Our work enabled us to identify motifs for uncharacterized alleles and is to date the predictor entirely trained on naturally presented peptides with the largest allelic coverage, including all frequent HLA-I alleles in the Caucasian population. However, the number of HLA-I alleles for which predictions are available (58 in total) is still smaller than what other tools can do (especially tools like NetMHCpan that can make predictions for any allele). Despite this limitation, we emphasize that our work provides the first scalable framework to integrate HLA peptidomics datasets that will be or are being generated for neo-antigen discovery in cancer immunotherapy and therefore will enable increasing the allele coverage as new studies are published. Moreover, our results suggest that improving existing models describing the binding specificity of common HLA-I alleles may be as important as expanding allele coverage to rare alleles for neo-antigen discovery.

All HLA peptidomics datasets used in this work were generated with only 1% FDR and are of high purity. For this reason, and also to prevent including potential biases or removing important data, we decided not to filter our data with existing HLA-I ligand predictors, but we expect some contaminants in our large sets of peptides. Moreover, in a few cases, the motifs for some alleles were not detectable (see HLA-C12:03 in Fig. 1). This suggests that the (few) peptides binding to this allele may contaminate the other motifs. This is a known situation when analysing HLA peptidomics data with unsupervised approaches [24,40,41]. As previously observed, it affects especially HLA-C alleles which are often poorly expressed and whose binding specificities are more redundant [24,42]. However, our results show that some level of noise is tolerated for training our predictor and can still lead to improvement over existing tools (Fig 4 and S9 Fig). Moreover, as both the throughput and the accuracy of HLA peptidomics technology keeps improving, we expect that motifs of HLA-C alleles will become increasingly detectable in future and larger datasets. We also stress that no existing HLA-ligand interaction dataset used for training predictors is free of false-positives and for many alleles the number of ligands used to train existing predictors is significantly lower than what was used in the predictor developed in this work (S5 Fig).

In a few cases, the motifs of two alleles could not be split because of the very high binding specificity similarity of these alleles (e.g., HLA-C07:01 and HLA-C07:02 in Fig 1B). We emphasize that this does not preclude the use of HLA peptidomics data for training HLA-I ligand predictors, since many other samples in our dataset contained only one of these two alleles together with other non-overlapping ones. As such our strategy takes advantage of our large collection of HLA peptidomics datasets to naturally overcome cases where the deconvolution could not be fully achieved in one given sample.

Direct identification of neo-antigens with MS shows higher specificity compared to our predictions based on exome sequencing data [15,17], as expected. However, it is important to realize that these experiments are challenging and can be carried out only in a small subset of patients with enough tumor material. Moreover, in many cases, no neo-antigen is found by MS. As such, our work provides a scalable approach to capitalize on large MS data obtained from some patients or cell lines in order to improve predictions of neo-antigens in other patients where MS analysis of the immunopeptidome could not be carried out and only exome sequencing data are available.

Previous computational approaches for neo-antigen predictions have shown that incorporation of gene expression data could improve accuracy [30,43]. Considering that our predictor using only exome sequencing information (i.e., the list of somatic mutations) already enabled us to improve predictions in the cancer samples analysed in this work, we anticipate that integrating additional information such as gene expression, when available, may lead to even more accurate predictions of neo-antigens or cancer testis antigens.

Results shown in Fig. 5 further emphasize the power of in-depth sampling of the HLA-I ligand space to inform us about molecular mechanisms underlying HLA-I binding properties [44]. Considering the rapid expansion of HLA peptidomics experiments performed in cancer immunotherapy research [16-18,20,26,27], we anticipate that our approach for HLA-I motif identification and annotation will enable similar analyses in the future to uncover other molecular determinants of HLA-I binding specificity.

Overall, our work shows for the first time that HLA-I motifs can be reliably identified across in-depth and accurate HLA peptidomics datasets without relying on HLA-I interaction prediction tools or *a priori* knowledge of HLA-I binding specificity. This unsupervised and scalable approach refines known HLA-I binding motifs and expands our understanding of HLA-I binding specificities to a few additional alleles without documented ligands. As such, this work is a powerful alternative to synthesizing every peptide for *in vitro* binding assays, or to genetically modifying [30] or transfecting cell lines with soluble HLA-I alleles [32,33,45]. Our results further contribute to our global understanding of HLA-I binding properties and improve neo-antigen predictions from exome sequencing data. This work may therefore facilitate identification of clinically relevant targets for cancer immunotherapy, especially when direct identification of neo-antigens with MS cannot be experimentally done.

## Methods

### Cell lines and antibodies

We carefully selected ten donors expressing a broad range of HLA-I alleles and generated novel HLA peptidomics data (Supplementary Table 1). EBV-transformed human B-cell lines CD165, GD149, PD42, CM467, RA957 and MD155 were maintained in RPMI 1640 + GlutaMAX medium (Gibco, Paisley, UK) supplemented with 10% FBS (Gibco) and 1% Penicillin/Streptomycin Solution (BioConcept, Allschwil, Switzerland). TIL were expanded from two melanoma tumors following established protocols [46,47]. Informed consent of the participants was obtained following requirements of the institutional review board (Ethics Commission, University Hospital of Lausanne (CHUV)). Briefly, fresh tumor samples were cut in small fragments and placed in 24-well plate containing RPMI CTS grade (Life Technologies Europe BV, Switzerland), 10% Human serum (Valley Biomedical, USA), 0.025 M HEPES (Life Technologies Europe BV, Switzerland), 55 μmol/L 2-Mercaptoethanol (Life Technologies Europe BV, Switzerland) and supplemented with a high concentration of IL-2 (Proleukin, 6,000 IU/mL, Novartis, Switzerland) for three to five weeks. Following this initial pre-REP, TIL were then expanded in using a REP approach. To do so, 25 × 10^6^ TIL were stimulated with irradiated feeder cells, anti-CD3 (OKT3, 30 ng/mL, Miltenyi biotech) and high dose IL-2 (3,000 IU/mL) for 14 days. The final cell product was washed and prepared using a cell harvester (LoVo, Fresenius Kabi). Leukapheresis samples (Apher1 and 6) were obtained from blood donors from the *Service régional vaudois de transfusion sanguine, Lausanne*. Upon receival of TIL and leukapheresis samples, the cells were washed with PBS on ice, aliquoted and stored as dry pellets at −80°C until use. High resolution 4-digit HLA-I typing was performed at the Laboratory of Diagnostics, Service of Immunology and Allergy, CHUV, Lausanne.

W6/32 monoclonal antibodies were purified from the supernatant of HB95 cells grown in CELLLine CL-1000 flasks (Sigma-Aldrich, Missouri, USA) using Protein-A Sepharose (Invitrogen, California, USA).

### Purification of HLA-I complexes

We extracted the HLA-I peptidome from 2-5 biological replicates per cell line or patient material. The cell counts ranged from 1 × 10^8^ to 3 × 10^8^ cells per replicate. Lysis was performed with 0.25% sodium deoxycholate (Sigma-Aldrich), 0.2 mM iodoacetamide (Sigma-Aldrich), 1 mM EDTA, 1:200 Protease Inhibitors Cocktail (Sigma, Missouri, USA), 1 mM Phenylmethylsulfonylfluoride (Roche, Mannheim, Germany), 1% octyl-beta-D glucopyranoside (Sigma) in PBS at 4°C for 1 hr. The lysates were cleared by centrifugation with a table-top centrifuge (Eppendorf Centrifuge 5430R, Schönenbuch, Switzerland) at 4°C at 14200 rpm for 20 min. Immuno-affinity purification was performed by passing the cleared lysates through Protein-A Sepharose covalently bound to W6-32 antibodies. Affinity columns were then washed with at least 6 column volumes of 150 mM NaCl and 20 mM Tris HCl (buffer A), 6 column volumes of 400 mM NaCl and 20 mM Tris HCl and lastly with another 6 column washes of buffer A. Finally, affinity columns were washed with at least 2 column volumes of 20 mM Tris HCl, pH 8. HLA-I complexes were eluted by addition of 1% trifluoroacetic acid (TFA, Merck, Darmstadt, Switzerland) for each sample.

### Purification and concentration of HLA-I peptides

HLA-I complexes with HLA-I peptides were loaded on Sep-Pak tC18 (Waters, Massachusetts, USA) cartridges which were pre-washed with 80% acetonitrile (ACN, Merck) in 0.1% TFA and 0.1 % TFA only. After loading, cartridges were washed twice with 0.1% TFA before separation and elution of HLA-I peptides from the more hydrophobic HLA-I heavy chains with 30 % ACN in 0.1 % TFA. The HLA-I peptides were dried using vacuum centrifugation (Eppendorf Concentrator Plus, Schönenbuch, Switzerland) and re-suspended in a final volume of 12 uL 0.1% TFA. For MS analysis, we injected 5 uL of these peptides per run.

### LC-MS/MS analysis of HLA-I peptides

Measurements of HLA-I peptidomics samples were acquired using the nanoflow UHPLC Easy nLC 1200 (Thermo Fisher Scientific, Germering, Germany) coupled online to a Q Exactive HF Orbitrap mass spectrometer (Thermo Fischer Scientific, Bremen, Germany) or with Dionex Ultimate RSLC3000 nanoLC (Thermo Fischer Scientific, *Sunnyvale, CA)* coupled online to an Orbitrap Fusion Mass Spectrometer (Thermo Fischer Scientific, San Jose, CA), both with a nanoelectrospray ion source. We packed an uncoated PicoTip 8μm tip opening with diameter of 50 cm × 75 um with a ReproSil-Pur C18 1.9 μm particles and 120 Å pore size resin (Dr. Maisch GmbH, Ammerbuch-Entringen, Germany) re-suspended in Methanol. The analytical column was heated to 50°C using a column oven. Peptides were eluted with a linear gradient of 2–30% buffer B (80% ACN and 0.1% formic acid) at a flow rate of 250 nl/min over 90 min.

Data was acquired with data-dependent “top10" method, which isolates the ten most intense ions and fragments them by higher-energy collisional dissociation (HCD) with a normalized collision energy of 27% and 32% for the Q Exactive HF and Fusion instruments, respectively. For the Q Exactive HF instrument the MS scan range was set to 300 to 1,650 *m/z* with a resolution of 60,000 (200 *m/z*) and a target value of 3e6 ions. The ten most intense ions were sequentially isolated and accumulated to an AGC target value of 1e5 with a maximum injection time of 120 ms and MS/MS resolution was 15,000 (200 *m/z*). For the Fusion, a resolution of 120,000 (200 *m/z*) and a target value of 4e5 ions were set. The ten most intense ions accumulated to an AGC target value of 1e5 with a maximum injection time of 120 ms and MS/MS resolution was 15,000 (200 *m/z*). The peptide match option was disabled. Dynamic exclusion of fragmented *m/z* values from further selection was set for 20 or 30 seconds with the Q Exactive HF and Fusion instruments, respectively.

### Data analysis of HLA-I peptides

We employed the MaxQuant computational proteomics platform [48] version 1.5.3.2 to search the peak lists against the UniProt databases (Human 85,919 entries, May 2014) and a file containing 247 frequently observed contaminants. N-terminal acetylation (42.010565 Da) and methionine oxidation (15.994915 Da) were set as variable modifications. The second peptide identification option in Andromeda was enabled. A false discovery rate of 0.01 was required for peptides and no protein false discovery rate was set. The enzyme specificity was set as unspecific. Possible sequence matches were restricted to 8 to 15 amino acids, a maximum peptides mass of 1,500 Da and a maximum charge state of three. The initial allowed mass deviation of the precursor ion was set to 6 ppm and the maximum fragment mass deviation was set to 20 ppm. We enabled the ‘match between runs’ option, which allows matching of identifications across different replicates of the same biological sample in a time window of 0.5 min and an initial alignment time window of 20 min.

### Publicly available HLA peptidomics data

To expand the number of samples and survey an even broader range of HLA-I alleles, we included in this study forty publicly available HLA peptidomics data from seven recent studies [17,18,22,23,25-27]. Only samples with HLA-I typing were used. Peptides identified in the recent study [18] in different repeats and under different treatments were pooled together to generate one list of unique peptides per sample. Since the published peptidomics datasets from Pearson et al. [27] were filtered to include only peptides with predicted affinity scores of less or equal to 1250 nM, we re-processed the mass spectrometer raw data using MaxQuant with similar settings as mentioned above except that peptide length was set to 8-25 mers (S2 Dataset). HLA typing information was retrieved from the original publications. In one case (THP-1 cell lines), the typing is controversial [49]. In this work, we used the typing determined by the authors of the HLA peptidomics study where the data came from [23]. The high fraction (>50% of the peptides) displaying a clear A24:02 and B35:03 motif based on our unsupervised deconvolution further indicates that these alleles are truly expressed in the sample on which the HLA peptidomics analysis was performed.

### IEDB data

Known HLA-I ligands were retrieved from IEDB (mhc_ligand_full.csv file) [4]. All ligands annotated as positives with a given HLA-I allele (i.e., “Positive-High", “Positive-Intermediate", “Positive-Low" and “Positive") were used to build the IEDB reference motifs (Fig. 2A and 2C). Ligands coming from HLA peptidomics studies analysed in this work were not considered to prevent circularity in the motif comparisons and because the HLA-I alleles to which these peptides bind were not experimentally determined. Position Weight matrices (PWMs) representing binding motifs in IEDB and used to compare with motifs derived from our deconvolution of HLA peptidomics datasets were built by computing the frequency of each amino acid at each position and using a random count of 1 for each amino acid at each position.

### HLA peptidomics data from mono-allelic cell lines

All 16 HLA peptidomics datasets obtained from mono-allelic cell lines were downloaded from [30]. Motifs used in the comparison presented in Fig 2B were built in the same way as for IEDB data. When benchmarking our ability to re-predict such data with, we considered 9-mers from the ten alleles that overlapped our set of deconvoluted HLA peptidomics data and added 99-fold excess of random peptides from the human proteome. The fraction of positives (i.e., MS peptides identified in these mono-allelic cell lines) predicted in the top 1% was used to assess the prediction accuracy with different methods (Given the large number of decoy, AUC values would all be larger than 0.99, and therefore not appropriate to compare HLA-I ligand predictors).

### Mixture model for HLA-I binding motifs identification in HLA peptidomics data

An algorithm based on mixture models and initially developed for multiple specificity analysis in peptide ligands [28,29] was used to identify binding motifs in each dataset analysed in this work. Briefly, all peptides pooled by mass spectrometry analysis of eluted peptide-HLA-I complexes in a given sample were first split into different groups according to their size (9-10 mers). All 9-and 10-mers ligands were then modelled using multiple PWMs [24]. The results of such analysis consist of a set of PWMs that describe distinct motifs for each HLA peptidomics datasets (see S1 Fig) and probabilities (i.e., responsibilities) for each peptide to be associated with each motif. The command-line script of this mixture model (“MixMHCp") can be downloaded at: https://github.com/GfellerLab/MixMHCp.

### Fully unsupervised deconvolution of HLA peptidomics datasets

The availability of high-quality HLA peptidomics data from several samples with diverse HLA-I alleles suggest that one could infer which motifs correspond to which HLA-I alleles without relying on comparison with known motifs. For instance, if two samples share exactly one HLA-I allele, it is expected that the shared motif will originate from the shared allele (Fig. 1). To exploit this type of patterns of shared HLA-I alleles, we designed the following algorithm:

1. For each allele present in at least two samples, find all samples that share this allele (e.g., HLA-A24:02 in Fig. 1). Identify the shared motif. If a shared motif is found, map this motif to the corresponding allele in each sample.
2. Find samples that share all except one allele with another sample (S2 Fig). Find the motif that is not shared and map it to the HLA-I allele that is not shared.
3. Check if some samples contain exactly one motif and one allele that have not yet been associated. Annotate the remaining motif to the corresponding allele.
4. Use the motifs mapped to HLA-I alleles in steps 1), 2) and 3) to identify them in other samples that contain these alleles based on motif similarity.
5. Go back to 1) until no new motif can be mapped to HLA-I alleles.

Comparison of motifs was performed using Euclidean distance between the corresponding PWMs: 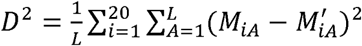, where *M* and *M’* stands for two PWMs (i.e., *20* × *L* matrices) to be compared and *L* is the peptide length. A threshold of T = 0.078 was used to define similar motifs based on visual inspection. Cases of inconsistencies (i.e., distances larger that T) between motifs mapped to the same allele were automatically eliminated. The final binding motif for each HLA-I allele (Fig. 2 and S4 Fig) was built by combining peptides from each sample that had been associated with the corresponding allele. Other measures of similarity between PWMs did not improve the results and we therefore used the Euclidean distance throughout this study.

In practice, HLA-A and HLA-B alleles tend to be more expressed and therefore give rise to a stronger signal in HLA peptidomics data. We therefore used first our deconvolution method [24], setting the number of motifs equal to the number of HLA-A and HLA-B alleles and identified HLA-A and HLA-B motifs with the algorithm introduced above (Step 1). We then ran our deconvolution method [24] without restricting the number of clusters (Step 2) and identified motifs corresponding to HLA-A and HLA-B alleles based on the similarity with those identified in Step 1. The remaining motifs were then analysed across all samples with the algorithm introduced above.

### Semi-supervised deconvolution of HLA peptidomics datasets

To expand the identification of binding motifs for alleles without known ligands, we used data from IEDB for HLA-I alleles with well-described binding motifs. In practice, for all HLA-I alleles in our samples that had not been mapped to motifs in the fully unsupervised approach and have more than twenty different ligands in IEDB, PWMs were built from IEDB data. These PWMs were used to scan the remaining motifs in each sample that contained the corresponding alleles. Motifs were mapped to HLA-I alleles if exactly one PWM obtained with the mixture model was found to be similar to the IEDB-derived motif (i.e., Euclidean distance smaller than T, as before). The unsupervised procedure described above was then applied to the remaining motifs to identify new motifs for alleles without ligands in IEDB.

### Amino acid frequencies at non-anchor positions in HLA peptidomics datasets

To have reliable estimates of the potential technical biases due to MS, amino acid frequencies were computed at non-anchor positions (P4 to P7) for alleles in our HLA peptidomics datasets (9-mers). Alleles showing some specificity at these positions (A02:01, A02:05, A02:06, A02:20, A25:01, A26:01, A29:02, B08:01, B14:01, B14:02, C03:03, and C07:04, see S4 Fig) were excluded from this analysis. The average frequencies of amino acids across alleles were then compared against the human proteome using Pearson correlation coefficient (Fig. 3). We also performed the same analysis with HLA-I ligands (9-mers) from IEDB splitting between those obtained by MS and those obtained by other assays (“non-MS data") (see S7 Fig). To enable meaningful comparison between these datasets, only alleles present in our HLA peptidomics data, with more than 100 ligands in both IEDB MS and non-MS data were considered in this analysis (14 alleles in total, see S4 Fig).

### Prediction of neo-antigens

For each HLA-I allele, PWMs were built from all peptides associated to this allele across all samples where the binding motif could be identified, using the highest responsibility values of the mixture model [24]. The frequency of each amino acid was first computed. Pseudocounts were added using the approach described in [50], based on the BLOSUM62 substitution matrix with parameter β=200. The score of a given peptide (*X*_*1*_,…*X*_*N*_) was computed by summing the logarithm of the corresponding PWM entries, including renormalization by expected amino acid frequencies: 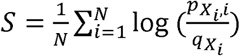. Here q_A_ stands for frequency of amino acid A at non-anchor positions (Fig. 3 and Supplementary Table 2), *p*_*A,i*_ stands for the PWM entry corresponding to amino acid *A* at position *i*, and *N* stands for the length of the peptide (*N*=9,10). The final score of a peptide was taken as the maximal score across all alleles present in a given sample and a P-value estimate was computed by comparing with distribution of scores obtained from 100,000 randomly selected peptides from the human proteome, so as to have similar amino acid frequencies compared to endogenous ligands.

To test our ability to predict neo-antigens, we used four melanoma samples in which ten neo-antigens (9-and 10-mers) have been directly identified with in-depth immunopeptidomics analyses of the tumor samples: Mel5, Mel8, Mel15 from [17] and 12T [20]. Missense mutations (i.e. cancer specific non-synonymous point mutations) identified by exome sequencing in those four melanoma samples [17] were retrieved and a list of all possible 9-and 10-mer peptides encompassing each mutation was built (S3 Dataset). Multiple transcripts corresponding to the same genes were merged so that each mutated peptide appears only once in the list. The total number of potential neo-antigens in each sample is shown in Table 1. Predictions for each HLA-I allele of each sample were carried out with the model described above. Peptides were ranked based on the highest score over the different alleles present in their sample. In parallel, affinity predictions with NetMHC (v4.0) [8] and NetMHCpan (v2.8) [12] and stability predictions with NetMHCstabpan (v1.0) [34] were performed for the same peptides and peptides were ranked based on predicted affinity using the highest value (i.e., lowest Kd) over all alleles. Only HLA-A and HLA-B alleles were considered since HLA-C alleles are known to show much lower expression and NetMHC could not be run for some HLA-C alleles in these melanoma patients. Ranking of the neo-antigens compared to all possible peptides containing a missense mutation is shown in Table 1 with either our predictor (“MixMHCpred") and the other tools mentioned above. Area Under the Curves were also computed (Fig. 4B), as well as the fraction of neo-antigens that fell among the top 1% of the predictions (Fig. 4C).

The same analysis was applied to study neo-antigens (9-and 10-mers) recently identified in two lung cancer patients [35] (L011: FAFQEYDSF, GTSAPRKKK, SVTNEFCLK, RSMRTVYGLF, GPEELGLPM and L013: YSNYYCGLRY, ALQSRLQAL, KVCCCQILL) and the ranking of these epitopes with different HLA-ligand predictors with respect to the full list of potential epitopes is shown in Supplementary Table 3 (see also Fig. 4B and 4C).

The standalone command-line programme “MixMHCpred" to run these predictions is provided free of charge for academic users (https://github.com/GfellerLab/MixMHCpred) to run HLA-ligand predictions with the models trained on HLA peptidomics data.

### Predictions of neo-antigens in melanoma samples – cross-patient validation

To assess how much our improved predictions of neo-antigens in the three melanoma samples of [17] depend on HLA peptidomics data generated from these samples, we performed a careful cross-sample validation. For each of the three samples where neo-eptiopes had been identified (Mel15, Mel8, Mel5) [17], we re-run our entire pipeline (i.e., annotation of HLA-I motifs across HLA peptidomics datasets + construction of PWMs for each allele) without the HLA peptidomics data coming from this sample. The PWMs were then used to rank all possible peptides (9-and 10-mers) encompassing each mutation. Overall, the predictions changed very little (S8 Fig).

### Predictions of cancer testis antigens

All cancer testis antigens with annotated epitopes (9-or 10-mers) and HLA restriction information were retrieved from the CTDatabase [36] for any of 58 alleles considered in our predictor (see Supplementary Table 4). All other possible peptides along these proteins (9-or 10-mers) were used as negatives when benchmarking the predictions on this dataset.

### Analysis of HLA sequences and structures

HLA-I sequences were retrieved from IMGT database [3]. All protein structures analysed in this work were downloaded from the PDB. Residues forming the P2 binding site in HLA-B14:02 (PDB: 3BVN [51]) were determined using a standard cut-off of 5Å from any heavy atoms of arginine at P2.

### Binding stability assays for HLA-B14:02 wt and W121R mutant

W121R mutation was introduced into HLA-B14:02 wt by overlap extension PCR and confirmed by DNA sequencing. BL21(DE3)pLys bacterial cells were used to produce HLA-B14:02 wt and W121R as inclusion bodies. Four peptides with histidine or arginine at P2 (A[H/R]TKPRPAL, G[H/R]YDRSKSL, A[H/R]FAKSISL, H[H/R]FEKAVTL) were synthesized at the Peptide Facility (UNIL, Lausanne) with free N and C-termini (1mg of each peptide, > 80% purity). Peptides with histidine at P2 come from our HLA peptidomics data and were assigned to HLA-B15:18 by our mixture model algorithm. Based on our analysis of HLA sequence and structure, peptides with histidine at P2 are predicted to interact with HLA-B14:02 W121R, while peptides with arginine at P2 are predicted to bind better HLA-B14:02 wt.

Synthetic peptides were incubated separately with denaturated HLA-B14:01 wt and HLA-B14:01 W121R mutant refolded by dilution in the presence of biotinylated beta-2 microglobulin proteins at temperature T=4°C for 48 hours. The solution was then incubated at 37°C and samples were retrieved at time t=0h, 8h, 24h, 48h and t=72h. The known HLA-B14:02 ligand IRHENRMVL was used for positive controls (measured half-live of 248h). Negative controls consist of absence of peptides. k_off_ were determined by fitting exponential curves to the light intensity values obtained by ELISA at different time points. Half-lives were computed as ln(2)/k_off_. Values shown in Fig. 5D correspond to the average over two replicates. For two peptides showing exceptionally high binding stability, only lower bounds on half-lives could be determined (dashed lines in Fig. 5D).

## Data availability

The mass spectrometry proteomics data have been deposited to the ProteomeXchange Consortium via the PRIDE partner repository with the dataset identifier PXD005231. The code to predict HLA-I ligands can be accessed at https://github.com/GfellerLab/MixMHCpred.

## Acknowledgment

We thank Nicholas MacGranahan and Charles Swanton for sharing with us the list of somatic mutations for the two lung cancer samples [35] and Mathieu Courcelles for sharing the HLA-C typing of cell lines used in ref. [27]. We are thankful to Camilla Jandus and Pedro Romero for sharing the B cell lines with us. We thank the Protein Analysis Facility of the University of Lausanne for technical help and access to MS instrumentation during the first period of this study. We thank Julien Racle and Santiago Carmona for insightful discussions about the manuscript.

## Author contributions

M.B.-S. contributed to the design of the study, performed HLA peptidomics experiments, analysed the data and wrote the manuscript. C.C. performed HLA peptidomics experiments. P.G. performed the mutagenesis and *in vitro* binding experiments. M.S. analysed the data. H.P. performed MS measurements. P.G. provided reagents. L.E.K provided reagents. G.C. provided reagents and contributed to the manuscript. D.G designed the study, developed and implemented the algorithms, analysed the data and wrote the manuscript.

